# Glacier retreat reshapes trophic networks: interaction turnover outpaces species turnover over space-time

**DOI:** 10.64898/2026.04.13.718237

**Authors:** Bao Ngan Tu, Eduard Mas-Carrió, Jiri Hodecek, Nora Khelidj, Morena Casartelli, Sergio Rasmann, Luca Fumagalli, Gianalberto Losapio

**Affiliations:** Department of Biosciences, University of Milan, 20133 Milan, Italy; Department of Ecology and Evolution, Laboratory for Conservation Biology, Biophore, University of Lausanne, Switzerland; Swiss Human Institute of Forensic Taphonomy, University Centre of Legal Medicine, Switzerland; Institute of Earth Surface Dynamics, University of Lausanne, Switzerland; Institute of Biology, University of Neuchatel, Switzerland

**Keywords:** Arthropod diet, Biodiversity change, Climate change, Food web assembly, Glacier extinction, Gut-content DNA metabarcoding, Plant–arthropod interactions, Trophic networks

## Abstract

As glaciers retreat worldwide, newly exposed terrains are rapidly colonized by plants and their associated animal communities. Although plant–animal interactions are key for biodiversity maintenance and ecosystem functioning, the ecological processes underlying the assembly and development of trophic interactions over space-time remain poorly understood. Here, we investigated the trophic niche structure of plant–arthropod interactions along a 140-year primary succession at Mont Miné glacier foreland (Switzerland). Using arthropod gut-content DNA metabarcoding, we reconstructed trophic interactions at the food web level, revealing numerous previously undetected links among 284 arthropod taxa feeding on 263 plant taxa. Trophic niche overlap among arthropods increased following glacier retreat, indicating decreasing resource partitioning and suggesting increased resource competition. Trophic niche breadth became narrower and diet species richness declined, indicating increased trophic specialization. Notably, changes in trophic interactions occurred more rapidly than shifts in species diversity or community composition of plants and arthropods. These results demonstrate that glacier retreat reorganizes trophic networks beyond simple species turnover, reshaping biotic interactions during ecosystem development. Our findings highlight trophic interactions as sensitive indicators of biodiversity change and suggest that the stability of emerging food webs may be strongly affected as glaciers vanish worldwide.

## INTRODUCTION

The retreat of mountain glaciers worldwide, caused by global warming, is particularly pronounced at low and medium latitudes such as at the European Alps (Zemp et al., 2015; Zemp et al., 2019; Roe et al., 2017). Current projections with both long-term (Rounce et al., 2023; Hock et al., 2019) or short-term model timescales (Cook et al., 2023) suggest that by 2100 large alpine glaciers could lose up to more than 60 % of their mass compared to 2015, while smaller glaciers would be completely gone (Raup et al., 2025). As of 2025, many glaciers have already gone extinct (Linsbauer et al., 2025), including Switzerland’s Pizol (Huss et al., 2025), Dolomites’ Fradusta Inferiore glacier (Securo et al., 2025), and Iceland’s Okjökull (Van Tricht et al., 2026). These rates of vanishing glaciers raise serious concerns about the potential complete disappearance of water reservoirs and the loss of irreplaceable, unique ecosystems and biomes. Indeed, glaciers play a critical role in mountain ecosystem functioning (Stibal et al. 2020; Casallas et al., 2025) by providing unique habitats to biodiversity as well as essential services to humans (Moore et al., 2009; Salim 2023). The loss of glacial systems therefore represents not only a reduction in meltwater availability but also a substantial threat to alpine biodiversity, including cold-adapted and often endemic plant and animal species of high conservation value (Cauvy-Fraunié and Dangles 2019; Wilkes et al., 2023; Bosson et al., 2023, Losapio et al., 2025a). Yet, the consequences of glacier loss for biotic interactions among plant and animal species remain poorly understood (Tu et al., 2024; Conti et al., 2024). Given that ecological interactions structure communities and sustain ecosystem functioning, understanding how glacier retreat reshapes interaction diversity is crucial for predicting biodiversity responses.

Following glacier retreat, newly exposed ice-free lands are rapidly colonized by living organisms, creating primary ecological succession and forming glacier foreland (Whittaker 1993; Bradley et al. 2014; Fickert et al., 2017; Eichel 2019; Fickert 2020). However, these unique ecosystems are highly sensitive to environmental change as, once lost, cannot be restored (Stibal et al., 2020; Goury et al., 2025). Previous studies have reported short-term increases in biodiversity following glacier retreat (Ficetola et al., 2024); however, this trend is often temporary (Ficetola et al., 2021, Losapio et al., 2021, Tu et al., 2024) as biodiversity declines in ‘climax’ communities at the end of the succession process (Poorter et al., 2023). Species diversity, functional diversity, and interaction network diversity commonly peak within 30–100 years after deglaciation depending on regional and climatic conditions (Boggs et al., 2010; Losapio et al. 2021; Anthelme et al. 2022; Nota et al., 2025). Over a timescale of approximately 100–150 years, these areas can develop into mature ecosystems, forming forested communities (Ficetola et al., 2021; Hanusch et al., 2022). While later successional stages may exhibit increased network complexity, they are often characterized by reduced biodiversity due to competition and replacement as communities approach climax or equilibrium (Tu et al., 2024; Conti et al., 2024; Losapio et al., 2025a, b; Yang et al., 2025). Recent advances emphasize the importance of biotic interactions in the change of ecosystem structure rather than focusing only on species composition and richness (Junker et al., 2020; Hanusch et al., 2022; Khelidj et al., 2026). In particular, trophic interactions and food webs are crucial for ecosystem functioning as they ensure the energy flow from primary producers to consumers (May 2009; Hågvar et al., 2024), ultimately influencing ecosystem stability and resilience (Pimm 1982; Dunne 2009). However, the ecological niche processes underlying on the assembly of these trophic interactions and food web dynamics remains poorly understood. This knowledge gaps hinder our ability to predict ecosystem dynamics, changes in carbon and nutrient cycling, and anticipating the impact of glacier extinctions on biodiversity.

Key aspects of food web structure can be captured by analysing the trophic niche, particularly: (1) the **diet species richness (DSR)**, defined as the number of consumed taxa in the diet of each consumer species, which represents a local measure of resource exploitation (Bolnick et al., 2003; Kiester, 2013; Roslin & Majaneva 2016; Diegle et al., 2019); (2) the **trophic niche overlap** (Pianka’s index), which quantifies the degree to which two consumer species share food resources within a given community and provides insights into potential niche partitioning or competition (Pianka 1973; Hulbert 1978); and (3) the **trophic niche breadth** (Levins’ index), which characterizes species along a specialist–generalist continuum (Cowell & Futuyma 1971; Feisinger et al., 1981). Understanding the response of these key indicators to climate change and particularly how glacier retreat reshapes trophic interaction networks is key for our knowledge of ecological stability.

Across diverse terrestrial and aquatic ecosystems, food web stability could be enhanced by high species richness, weak interaction strength, and high trophic niche overlap by promoting functional redundancy and by providing alternative pathways for energy flow (Dunne et al., 2002; Quevedo et al., 2009; Gellner et al., 2023; López-Rodríguez et al., 2025). In proglacial lakes, food web complexity and biodiversity increase following lake age (>150 years) (Tiberti et al., 2020; Jeppesen et al., 2023; López-Rodríguez et al., 2025). Early successional ecosystems, such as new ice-free terrains, are typically characterized by structurally simple food webs dominated by generalist consumers and high niche overlap (Piechnik et al. 2008). As succession proceeds and resource diversity increases, specialist species become more prevalent and trophic niche overlap decreases as resource partitioning and niche differentiation promote the stability of food webs (Hulbert 1978, Quevedo et al., 2009). Evidence from proglacial systems suggests that pioneer communities, such as those involving Collembola, exhibit highly flexible trophic interactions during early succession (Hågvar et al., 2020). Despite growing evidence on food webs in proglacial ecosystems, little is known about how trophic niches of plants and arthropods develop through space and time as glaciers retreat. Understanding changes in trophic interaction structure following glacier retreat would provide novel insights into how interactions between primary producers (plants) and primary consumers (phytophagous arthropod) originate, assemble, and develop over spacetime.

A possible limitation to the advancement of trophic interaction studies is that food web analysis in terrestrial ecosystems usually relies on traditional observational methods that capture only few interactions for a restricted number of animals and taxa (Roslin & Majaneva 2016; Diegle et al., 2019) while do not fully reflect the diversity and complexity of producer–consumer interactions at ecosystem scale (Symondson 2002). To overcome these limitations, this study uses molecular ecological network analysis (Meyer et al., 2020), particularly organismal DNA metabarcoding to examine arthropod gut contents and identify arthropod diet, a powerful approach that allows us to explore non-observable trophic interactions of ‘who eats whom’ and enables the detection of multiple trophic interactions from individual samples at once (Matheson et al., 2008; García-Robledo et al., 2013; Roslin et al., 2019; Pitteloud et al., 2023; Mas-Carrió et al., 2024). The combination of this approach within ecological succession and niche theory ultimately provides a comprehensive understanding of the processes underlying plant–arthropod trophic interaction network dynamics.

Our study addresses this shortcoming by examining how the trophic niche of plant–arthropod interactions change along a 140-year glacier retreat chronosequence. By integrating gut-content DNA metabarcoding with network analysis, we aim to uncover how trophic interactions and resource partitioning respond to glacier retreat and to changes in community structure and composition. Our novel approach allows us to assess how glacier retreat reshapes trophic interaction networks through changes in resource diversity, consumer specialization, and trophic niche partitioning. Specifically, we addressed three main research questions and tested the following hypotheses: 1) How do arthropods’ diet species richness and local diet change following glacier retreat? We hypothesise that diet species richness would increase towards mid-succession and then decline as plant diversity decreases because woody species replace diverse herbaceous communities in late succession; we also hypothesise that the proportion of local plant resources consumed by arthropods (hereafter ‘local diet’) will increase with increasing host plant abundance following glacier retreat. 2) How do arthropods’ diet similarity change following glacier retreat? We hypothesise that the similarity of trophic interactions will increase as plant species diversity decreasing in later deglaciation stage. 3) How do arthropods’ trophic niche breadth and trophic niche overlap change following glacier retreat? We hypothesise a shift toward more specialist feeding strategies as reflected by a decline of trophic niche breadth, we also hypothesise that trophic niche overlap is initially high at the early stage, and decreases as plant diversity increases; ultimately, at late stages, we expect that resource limitation would force multiple consumer species to exploit the same, few dominant plant species.

## MATERIALS AND METHODS

### Study area

Our study was performed along the Mont Miné glacier foreland in Valais, Switzerland (46°03′33.646′N, 07°32′54.550′E). Following recent global warming, Mont Miné glacier has lost 2.53 km in length since the end of the Little Ice Age (*c* 1850) (Nicolussi et al., 2022). Using evidence from existing geochronology of Mont Miné glacier (Lambiel et al., 2016; Nicolussi et al., 2022), complemented with field validation of moraine margins (Tu et al., 2024), we divided the Mont Miné foreland into four deglaciation stages delimited by moraines deposited in the following years: 1989, 1925, 1900, and 1864 (Fig. 1). Accordingly, we delimited Stage 1 between glacier tongue in 2022 and 1989, Stage 2 between 1989–1925, Stage 3 between 1925–1900, and stage 4 between 1900–1864. We then calculated the average terrain age as the mean years since glacier retreat between two adjacent moraines, resulting 17 years for Stage 1, 65 for Stage 2, 110 years for Stage 3, and 140 years for Stage 4. Following glacier dynamics, the ecosystem development ranges from patchy (Stage 1) and pioneer grasslands (Stage 2) to shrubland (Stage 3) and ‘climax’, closed forests (Stage 4) (for more details on site conditions, see Delarze et al., 2015; Price et al., 2021; Tu et al., 2024).

**Figure 1.**
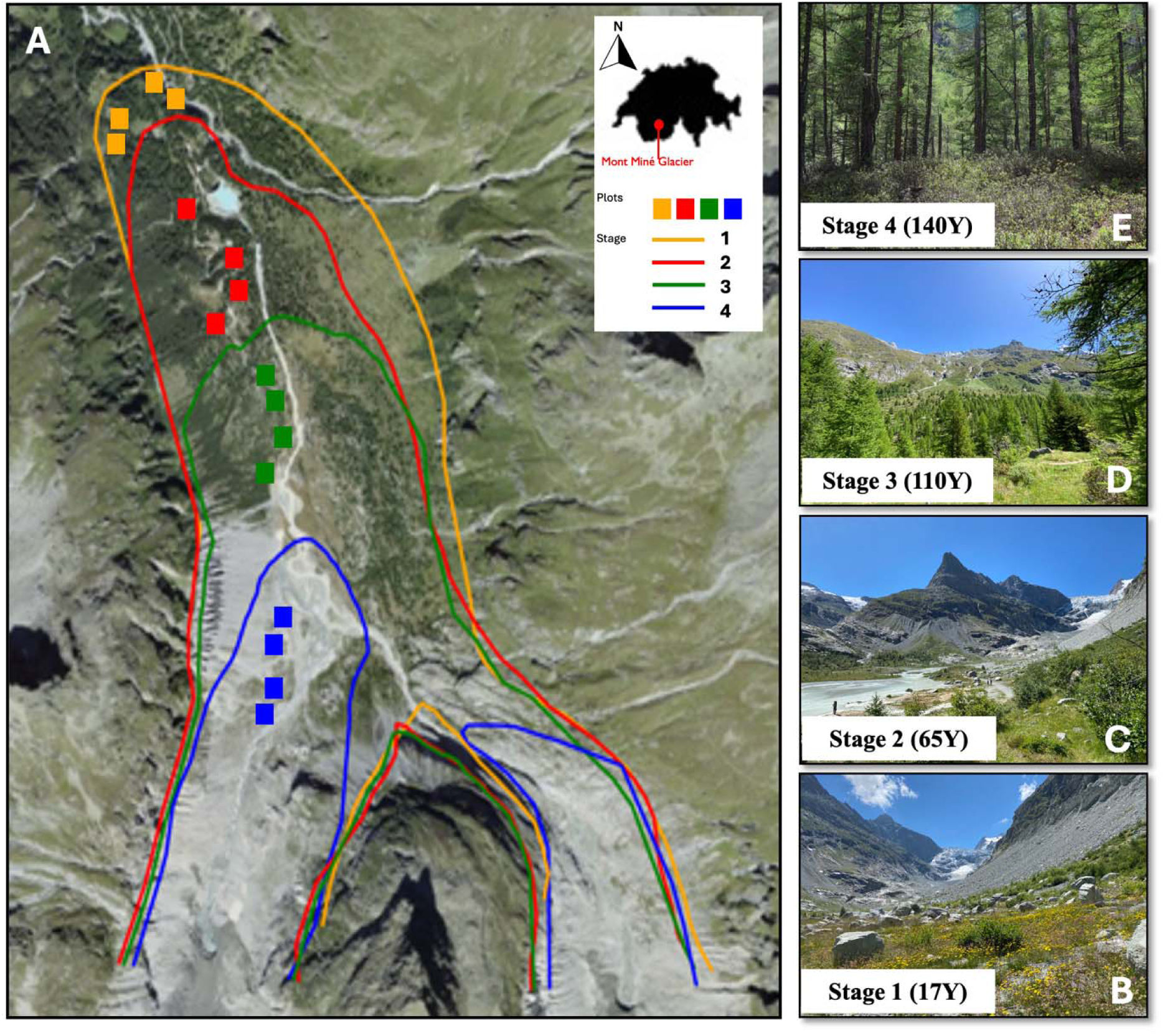
Study area: (A) Mont Miné glacier foreland; (B, C, D, E) Habitat pictures at deglaciation stages (Y = terrain age as years since glacier retreat): Stage 1 = 17 Years, Stage 2 = 65 Years, Stage 3 = 110 Years, Stage 4 = 140 Years.

### Sample collection and specimen identification

The study was conducted during summer 2022 (June and July), following the plant growing and blooming season, on dry and sunny days. In each deglaciation stage, we randomly set up n = 4 different plots (*n* = 16 total) of 3 x 3 m each. The distance between plots ranged from ca. 35m (closest: plot 1A to plot 1B) to 1830m (farthest: plot 1D to plot 4A) with the elevation ranges from 1961 to 2000 m a.s.l.

Plant communities were surveyed in each plot by identifying plant species and recording their coverage (approximation of 5%). Nomenclature follows Flora Helvetica (Lauber & Wagner, 1996; https://www.infoflora.ch); all plants were identified at species level for a total of n = 130 plant species belonging to 32 families (Table S1). Early-succession stages (Stage 1 and Stage 2) included pioneer species which are mostly or exclusively found in pioneer stages, such as *Linaria alpina, Oxyria digyna, Saxifraga aizoides,* and *Hieracium staticifolium*. In the mid-succession stages (Stage 3), more competitive species as well as woody species become dominant such as *Achillea erba-rotta, Trifolium pallescens, Juncus trifidus, Calluna vulgaris,* and *Alnus viridis*. As the whole Mont Miné glacier foreland lies in the subalpine zone, Stage 4 was characterized by climax-distinctive coniferous-forest species such as *Larix decidua*, *Picea abies*, *Rhododendron ferrugineum* and *Vaccinium myrtillus.* Generalist species, which occur in the whole glacial foreland, include *Anthoxanthum alpinum, Poa alpina, Leontodon helvesticus, and Lotus corniculatus* (Tu et al., 2024). Following vegetation survey, fresh and clean leaves were collected for building the local reference library.

To study arthropod communities and trophic interactions, we used two complementary methods for collecting samples. For arthropods visiting flowers and standing on leaves, we actively collected individual arthropods using swept net (Gibson et al. 2011; Grange et al. 2021). Sampling was carried out for each plot for a standard amount of observation time equal to 30 minutes per sampling round. Plots were sampled in random order of space (deglaciation stage) and time of the day. We conducted n = 6 temporal replicates throughout the blooming season per each plot, for a total of n = 96 sampling rounds. For ground-dwelling arthropods, we used wet pitfall traps (Knapp & Ruzicka 2012; Nakamura et al., 2020). One pitfall trap was set up for each plot, filled by propylene glycol and was left in use for 5 days in the field. Arthropods were preserved in Ethanol 90% initially at -20^°^C, then at -80^°^C, prior to DNA extraction. We performed n = 5 rounds of pitfall sampling during the sampling season (n = 80) (a final round of sampling was not possible due to the August heatwave).

Arthropod samples were then hand-sorted, photographed and identified to highest feasible taxonomic level. All the pictures and information were also shared using a citizen science project on the *iNaturalist* platform to gather further taxonomic information, for data sharing, and for public outreach (https://www.inaturalist.org/projects/pollinator-diversity-at-ferpecle-glacier-ecosystems). For complementary taxonomic ID of the morphologically indistinguishable arthropods, we run Sanger sequencing for a subset of n = 570 specimens. Trophic guild for each arthropod taxa was determined using diet coding from Rainford and Mayhew (2015) (Rainford and Mayhew 2015), including carnivore, detritivore, herbivore, omnivore, pollinator, pollinator and detritivore (for unclassified arthropods). Then, in total, we analyzed the diet of n = 919 arthropod samples belonging to 87 Families and 8 Orders (Supplement - Dataset).

### DNA extraction

For local reference library (plant samples), DNA was extracted from dried leaves using the Qiagen 96 Dneasy Plant DNA kit according to the manufactures’ instructions.

For gut-content DNA metabarcoding (arthropod samples), all the steps were done for each individual (sample) following the protocol of Koester & Gergs 2017 and Wallinger et al., 2013 with some adaptations. To reduce the amplification of DNA from plant materials sticking on the outer surface of the arthropods, we washed the sample first with sterile water to remove the ethanol, second with sodium hypochlorite (‘bleach’, 1–1.5%, 30s), then rinsed twice with sterile water. We dried and dissected to collect only the abdomens. For the small arthropods (∼ 1mm length), we keep the entire body and placed all individuals (of same species * sampling replicate * plot, eg. in case of Collembola, max 12 individuals) into one sample to maximize the plant DNA extracted. Samples were kept on liquid nitrogen and were homogenized with 0.5mm ceramic beads for 1 min at 6500 rpm using a MagNA Lyser Instrument in cold room. We added 200μL digestion mix (180μL ATL + 20μL proK) to each Eppendorf and incubated at 56^°^C 150 rpm overnight. Next steps of extractions were performed using the Blood and Tissue Kit (QIAGEN, USA) following the manufacturer protocol. All DNA samples were diluted 5-fold prior to PCR amplification to avoid the inhibitor (Mas-Carrió et al., 2024). All extractions were performed in a laboratory restricted to low DNA-content analyses (LBC lab, DEE, UNIL, Lausanne, Switzerland).

### Sanger sequencing

For local reference library, plant DNA extracts were amplified using DNA barcoding of the P6 loop of the trnL intron, chloroplast DNA (primer Sper01), targeting all vascular plant taxon (Spermatophyta) (Taberlet et al., 2018; Mas-Carrió et al. 2024). For complementary taxonomic of arthropods, arthropod DNA (from gut-content DNA extracts) were amplified using DNA barcoding of the COI region (Hodecek et al., 2024). PCR cycling programs were followed the protocol with some modifications (see Supplement for detailed laboratory procedure). PCR products were sent for Sanger sequencing in both directions at Microsynth AG (Balgach, Switzerland). The obtained SPER01 and COI sequences were aligned and blasted in GenBank, to relate them to reference sequences and identify them.

### Gut-content DNA metabarcoding

The plant chloroplast DNA (P6 loop of trnL intron) from gut-content DNA extracts were amplified separately for each sample in three replicates, using Sper01 primer and following the protocol and thermal cycling programs (Supplement). Each PCR plate contained 12 blanks, 4 extraction control, 6 negative and 1 positive PCR controls (DNA assembly of 2 plant species with increasing relative concentrations) and 73 DNA extracts (samples).

PCR products were run on a 1.5% agarose gel, stained with ethidium bromide and then visualized under UV light to confirm the amplification success and fragment sizes (fragment analysis was done at Lausanne Genomic Technologies Facility, UNIL). Then, PCR products were subsequently pooled per PCR plate and purified using a MinElute PCR Purification Kit (Qiagen, Hilden, Germany). Purified pools were quantified using a Qubit® 2.0 Fluorometer (Life Technology Corporation, USA). Library preparation was done following the TagSteady Protocol (Carøe and Bohmann, 2020). After adapter ligation, libraries were validated on a fragment analyzer (Advanced Analytical Technologies, USA). Final libraries were quantified, normalized and pooled before 150-bp paired-end sequencing on an Illumina MiniSeq sequencing system with a High Output Kit (Illumina, San Diego, CA, USA). These steps were done in LBC lab.

### Bioinformatic data analyses

The bioinformatic processing of the raw sequence output was performed using the OBITools package (Boyer et al., 2016) and was followed all the steps as in Mas-Carrió et al. (2024) with the following modifications (Supplement). Initially, forward and reverse reads were assembled with a minimum quality score of 40. The joined sequences were assigned to samples based on unique tags combinations. Assigned sequences were then de-replicated, retaining only unique sequences. All sequences with less than 20 reads per library were discarded as well as those not fitting the range of metabarcode lengths. Remaining sequences were assigned to taxa using a reference database for Sper01 generated from the EMBL database (European Molecular Biology Laboratory) with additional sequences from our local plant reference (as mentioned above). Further data cleaning and filtering was done in R (version 4.4.1) using the metabaR package (Zinger et al., 2021). Sequences that were more abundant in extraction and PCR controls than in samples were considered as contamination and removed.

Operational taxonomic units (OTUs) with similarity to the reference sequence lower than 90 % were also eliminated from the dataset. Removal of tag-leaked sequences was done independently for each library. Remaining PCR replicates were merged by each sample, keeping the mean relative read abundance (RRA), frequency of occurrence (FOO) and presence–absence (Mas-Carrió et al., 2024). Plant taxa in the diet of arthropod were determined at the highest level, if possible.

### Statistical analysis and modelling

All downstream analyses were carried out using R software (Version 4.4.1).

To have an overview of the arthropod community’s diet in the different deglaciation stages we calculated the dissimilarity matrix (Bray-Curtis distance) for each arthropod sample based on the final table of RRA, at plant species level, and examined the variation in plant-based diet composition between arthropod individuals using Principal Coordinate Analysis (PCoA). We visually illustrated the most abundant plant families of arthropod dietary composition.

To answer the first question “How do arthropods’ diet species richness and local diet change following glacier retreat?”, we calculated diet species richness (DSR) of each arthropod species across each guild within each plot as the number of distinct plant species detected in its diet using presence–absence data (Kiester, 2013). Duplicated trophic interactions within a plot were removed to ensure that each plant species was counted only once per arthropod species. This metric represents the taxonomic richness of resource use, with higher values indicating broader diets. Then, for local diet (plant species presented in both diet and plot), we computed the read relative abundance of those species in arthropods’ diet to examine shifts in local dietary tracking.

To answer the second question “How do arthropods’ diet similarity change following glacier retreat?”, precisely, to study the homogeneity of arthropod diet within one stage and to test whether the dietary composition differed significantly between stages, we computed Beta diversity (using Sorensen similarity) of Plant community composition, Arthropod community composition and Arthropod diet composition (species level) between plots within stage. The higher the index, the more similar the composition between plots within stage.

To answer the third question “How do arthropods’ trophic niche breadth and trophic niche overlap change following glacier retreat?”, we calculated trophic niche breadth of each arthropod species across guilds and deglaciation stages using Standardized Levins’ index, based on presence–absence data of consumed plant taxa (Equation number 1, 2; Levins 1968):

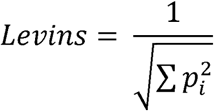

where *p_i_* represents the proportional use of resource *i*. Because presence–absence data were used, proportional use was calculated assuming equal weights across all detected plant taxa. To allow comparison among species with differing numbers of available resources, Levins’ index was standardized as:

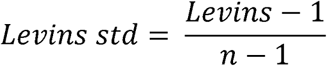

where *n* is the number of plant taxa considered. The standardized index ranges from 0 (extreme specialization) to 1 (maximum generalization). Species with no detected plant associations were excluded from analyses. Then, we computed trophic niche overlap among arthropod species across guilds and deglaciation stages using Pianka’s index (Equation number 3, Pianka 1973):

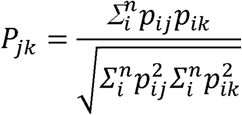

where *p_ij_* and *p_ik_* are the proportional uses of resource i by species j and k, respectively, and n is the total number of plant species. An overlap index (P_jk_) ranges from 0 (no overlap) to 1 (complete overlap), with values of 0–0.29 indicating no overlap, 0.30–0.59 moderate overlap, and ≥ 0.60 high overlap between species (modified from Grossman, 1986; Wei et al., 2025). We used the presence–absence dataset.

Finally, to estimate the effects of deglaciation stage (years since glacier retreat), plant richness and arthropod trophic guilds on dietary indices, we fitted generalized linear mixed models (GLMM) with these effects as fixed effects, separately and interaction, and plot, taxa and guild as random effects on the dietary data (Chambers and Hastie, 1992; Brooks et al., 2017), using the *glmmTMB* R package (Brooks et al., 2017), as follow:

_ Model 1: Diet_species_richness ∼ stage * plant_richness * guild + (1 | plot)

_ Model 2: Local_diet ∼ stage + plant_richness + guild + (1 | plot)

_ Model 3: Arthropod_diet_similarity ∼ stage + (1 | plot1) + (1 | plot2)

_ Model 4: Plant_community_similarity ∼ stage + (1 | plot1) + (1 | plot2)

_ Model 5: Arthropod_community_similarity ∼ stage + (1 | plot1) + (1 | plot2)

_ Model 6: Trophic_niche_breadth ∼ stage + guild + (1 | arthropod_taxa)

_ Model 7: Trophic_niche_overlap ∼ stage + (1 | guild) + (1 | arthropod_taxa1) + (1 | arthropod_taxa2)

The model 1 were fitted with using a negative binomial distribution with a log link function (nbinom2). The models from 2 to 7 were fitted with a beta distribution, zero inflation = ∼ 1, and heterogeneous dispersion among stages and guilds. Data distribution was assessed using the *summary* function (Chambers and Hastie, 1992). We report results in terms of explained variance by each predictor and estimated model parameters for each factor.

## RESULTS

### Gut-content DNA metabarcoding reveals cryptic arthropod food consumption

After quality filtering, we retained 4,676,768 reads of 1,243 different OTUs (Operational Taxonomic Units, or unique sequences) for the Sper01 assay that were assigned to 263 plant taxa (52 % identified to species level, 77.5 % to genus level, 99 % to family level, and 100% to order). Most frequent taxa were Asteraceae (694,275 reads), *Rhododendron ferrugineum* (383,481 reads), *Salix sp.* (328,172), *Lotus corniculatus* (295,142), Poaceae (214,992), which represent more than 40% of the sequences retained (Supplement - dataset).

Diet composition of arthropod guilds shifted across deglaciation stages (Figure 2) as PCoA ordination revealed deglaciation stages characterizing discrete clustering in diet composition (Figure S1). Arthropod diet composition differed significantly among deglaciation stages (PERMANOVA: F = 10.92, R² = 0.035, p < 0.001, Table S1). Furthermore, pairwise PERMANOVA revealed significant differences between all combinations of contrasts between stages (all p < 0.001, Table S1), with the highest differences reported between the earliest (stage 1) and latest (stage 4) stages (F = 22.67, R² = 0.054, Table S1), and the smallest between stages 2 and 3 (F = 5.23, R² = 0.010, Table S1). Multivariate dispersion also significantly differed among stages (PERMDISP: F = 8.12, p < 0.001, Table S1), indicating that some stages have more diverse diets than others. Vectors alignment indicated driving plant taxa (four most strongly correlated) for a particular deglaciation stage were overlaid on the ordination. The vector for Asteraceae was oriented toward stage 1 centroids, while the vector for *Pinus* (Pinaceae) showed a stronger association with stage 4 (Figure S1). The most frequently consumed plant families were Poaceae, Ericaceae, Asteraceae, Pinaceae, Fabaceae, Salicaceae, Betulaceae, and Campanulaceae, although their relative contributions varied among trophic guilds and deglaciation stages (Figure 2). Asteraceae and Fabaceae were particularly prominent in arthropod diets during the earliest glaciation stage (stage 1), whereas Ericaceae contributed more strongly to arthropod diets in the latest stage (stage 4).

**Figure 2.**
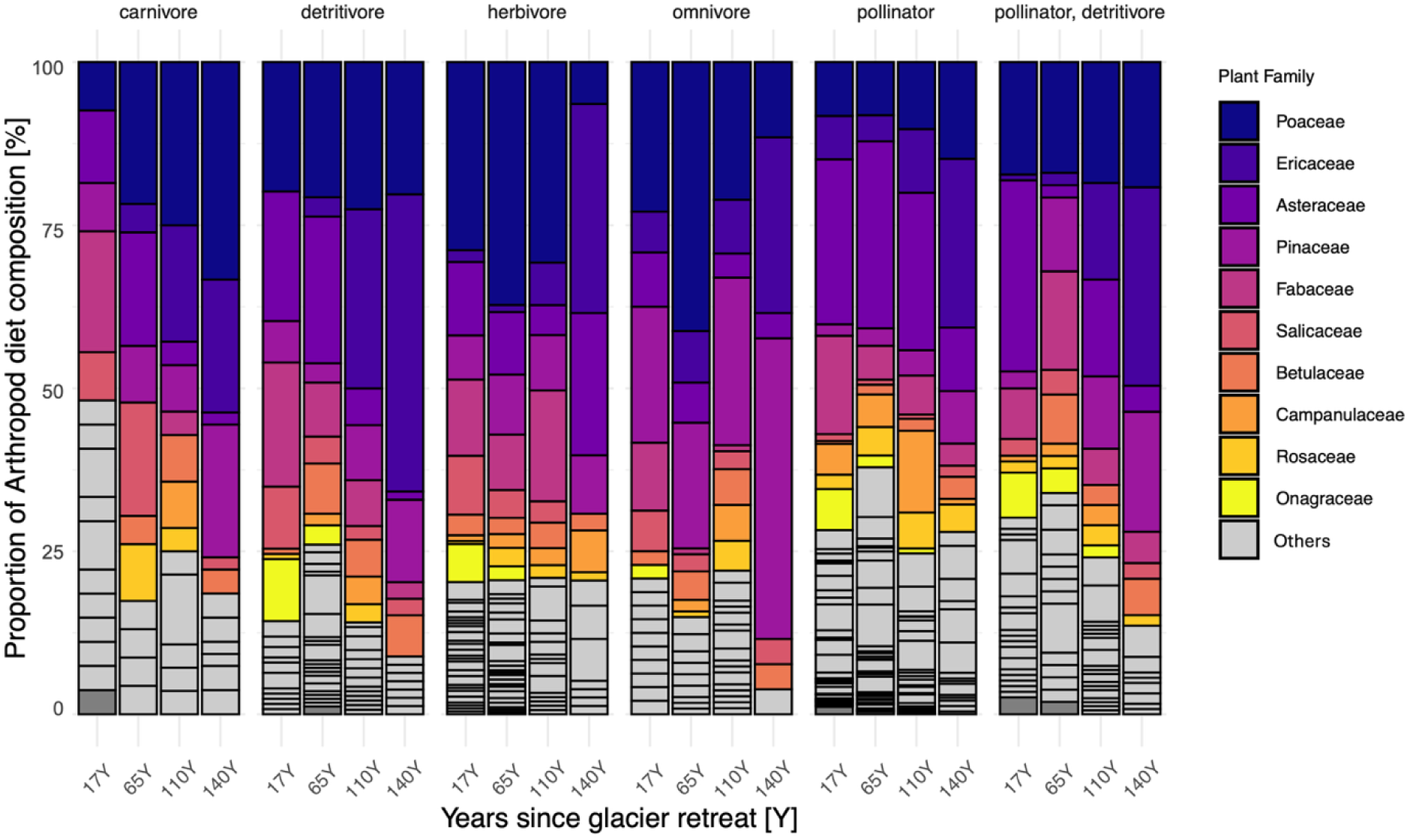
Stacked bar plots showing average percentage of arthropod diet composition (at plant family level, for clarity) across trophic guilds and among deglaciation stages (17, 65, 110, and 140 years since glacier retreat). Colored side bars indicate the most abundance plant families consumed.

### Diet species richness and local diet

DSR exhibited a non-linear response to glacier retreat (χ² = 7.95, p = 0.047, Model 1 - Table S1, Figure 3A). In the earlier stages, DSR initially increased as the highest rate was at intermediate stages, approximately 65 years since glacier retreat, while ultimately decreased at the oldest stages (∼140 years since glacier retreat). DSR significantly varied among arthropod trophic guilds (χ² = 116.46, p < 0.001, Model 1 - Table S1) depending on deglaciation stage (stage × guild interaction: χ² = 43.89, p < 0.001, Model 1- Table S1). Interestingly, plant species richness alone did not have a significant main effect on DSR (χ² = 0.04, p = 0.84, Model 1 - Table S1); however, its influence depended on deglaciation stage (stage × plant richness: χ² = 10.96, p = 0.012, Model 1 - Table S1). Although the three-way interaction among deglaciation stage, plant richness, and trophic guild was not significant (χ² = 13.37, p = 0.57, Model 1 - Table S1), a plant richness × guild interaction (χ² = 10.45, p = 0.064, Model 1 - Table S1) indicated that plant diversity may differentially influence dietary richness across feeding strategies.

**Figure 3.**
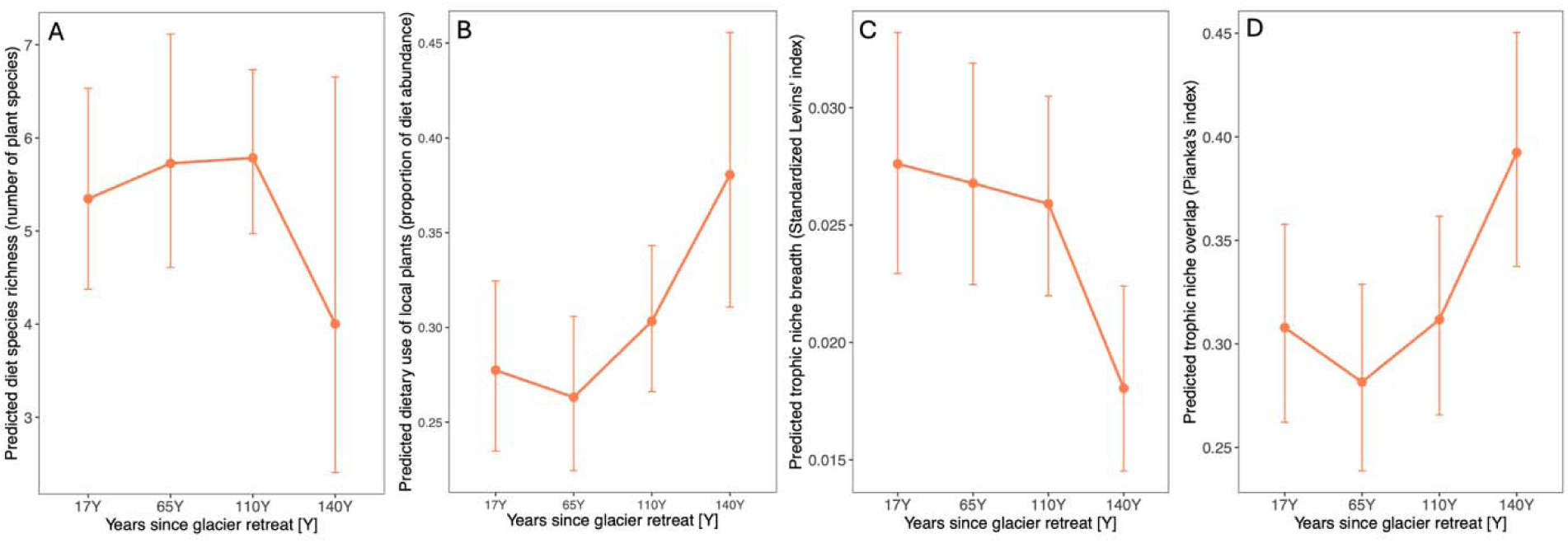
Predicted dietary metrics of arthropod communities across deglaciation stages. Estimated marginal means from generalized linear mixed models (glmmTMB) showing: (A) Predicted diet species richness (number of plant species consumed); (B) Predicted dietary use of locally available plants (proportion of diet abundance); (C) Predicted trophic niche breadth (Standardized Levins’ index); and (D) Predicted trophic niche overlap (Pianka’s index) across four deglaciation stages (17, 65, 110, and 140 years since glacier retreat). Points represent back-transformed marginal means and error bars indicate 95% asymptotic confidence intervals.

Accounting for plant taxa detected both in arthropod diets and within local plots, we found that the use of local food resources significantly increased following glacier retreat (χ² = 8.55, p = 0.036, Model 2 - Table S1, Figure 3B). Particularly, the proportion of local available plants in arthropod diets increased the most in the latest successional stage compared to early stages (stage 4: β = 0.47, p = 0.014, Model 2 - Table S2). Additionally, the presence of a plant species in the plot also strongly increased its probability of being detected in arthropod diets, indicating that the higher the abundance, the higher the consumption (χ² = 4.70, p = 0.030, Model 2 - Table S1). No differences were detected among trophic guilds (χ² = 10.13, p = 0.07, Model 2 - Table S1) as the positive relationship between deglaciation stage and local resource use was consistent across all guilds.

### Beta diversity

Notably, Beta diversity (Sorensen similarity) of arthropod dietary composition exhibited a strong and highly significant stage-dependent divergence, with within-stage dietary similarity declining sharply from early to late succession (χ² = 36.32, p < 0.001, Model 3 - Table S1, Figure 4A). Dietary dissimilarity was already evident at intermediate stages (stage 3: β = −0.30, p = 0.005, Model 3 - Table S1) and became most pronounced in the latest stage (stage 4: β = −0.62, p < 0.001, Model 3 - Table S1). We found a detectable effect of stage on the similarity of plant species composition within stage, with the highest similarity in early deglaciation stage (stage 1), a significant increase in dissimilarity towards stage 3 (β = −0.51, p = 0.021, Model 4 - Table S1), and a final increase in similarity at stage 4 (β = −0.07, p = 0.76, Model 4 - Table S1, Figure 4B). In contrast, the heterogeneity of arthropod community composition within stage consistently increased following glacier retreat, with significantly lower within-stage similarity across all later stages relative to the earliest stage (χ² = 11.62, p = 0.009, Model 5 - Table S1, Figure 4C).

**Figure 4.**
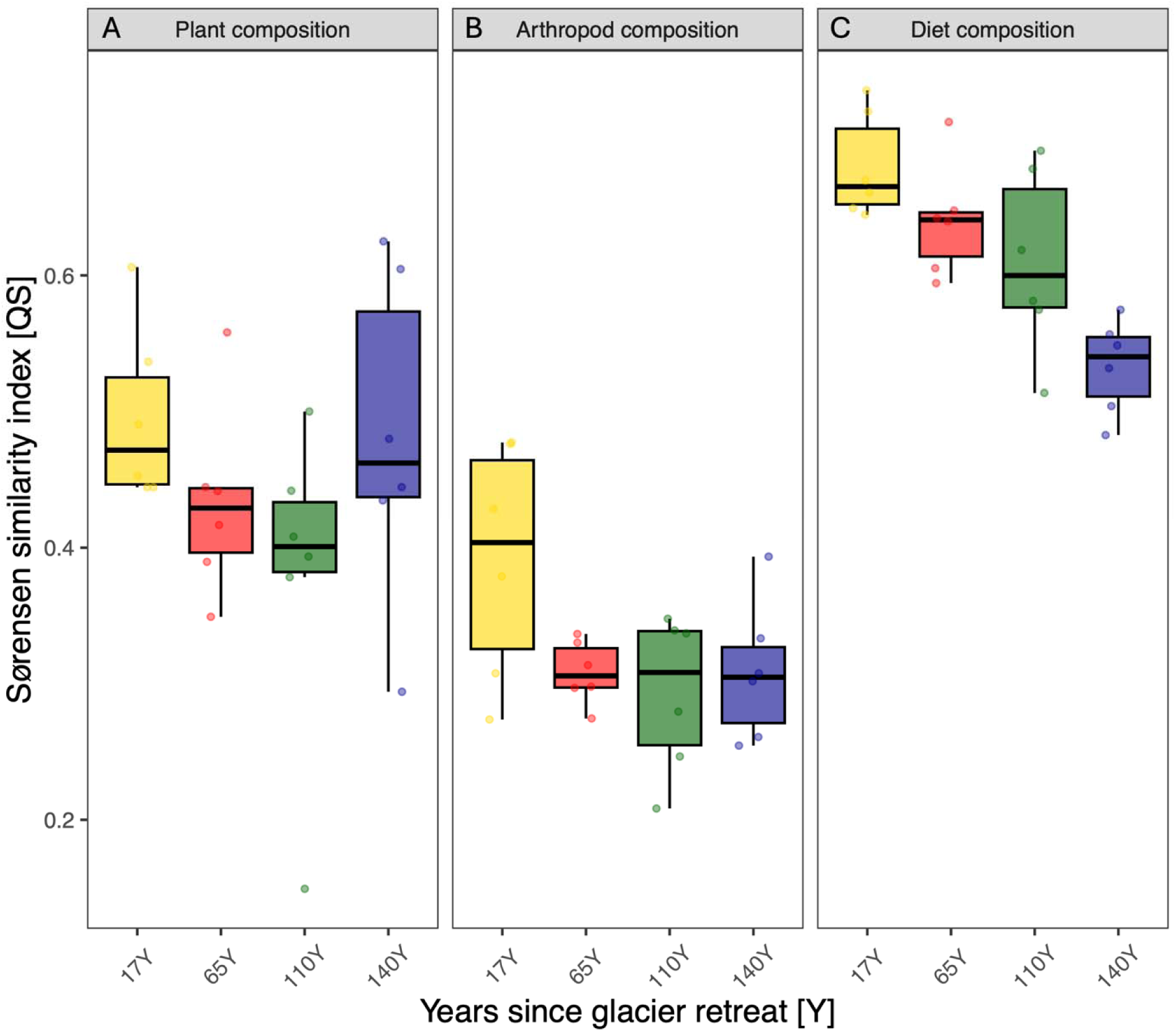
Compositional similarity, calculated by means of Sørensen index (QS), among different plots within deglaciation stage (4 plots per stage) (17, 65, 110, and 140 years since glacier retreat): (A) Similarity in the diet among arthropod species (∼ 263 plant taxa in arthropod diet); (B) Similarity in plant species composition at community level (∼ 130 plant taxa); (C) Similarity in arthropod species composition at community level (∼ 284 arthropod taxa).

### Trophic niche breadth

We found that trophic niche breadth (Standardized Levins’ index) changed significantly among deglaciation stages and trophic guilds (stage: χ² = 12.9, p = 0.0049; guild: χ² = 32.5, p < 0.001, Model 6 - Table S1, Figure 3C). Trophic niche breadth declined significantly in the latest stage relative to the earliest stage (stage 4: β = −0.43, p < 0.001, Model 6 - Table S1), indicating the increase of dietary specialization as succession ended. In contrast, we did not observe any significant differences in trophic niche breadth among intermediate stages (Model 6 - Table S1). We also found evidence for differences in trophic niche breadth among trophic guilds as pollinators exhibited significantly broader diets than both herbivores and omnivores (herbivore − pollinator: β = −0.49, p < 0.001; omnivore − pollinator: β = −0.69, p < 0.001; Pairwise Comparisons of Estimated Marginal Means, Model 6 - Table S1), whereas no significant differences in niche breadth were detected among the remaining guilds (all pairwise p > 0.10) (Model 6 - Table S1).

### Trophic niche overlap

We observed a significant increase of trophic niche overlap (Pianka’s index) between arthropod species following glacier retreat (χ² = 217.9, p < 0.001, Model 7 - Table S1, Figure 3D). Relative to the earliest stage (stage 1), we observed that mean trophic niche overlap among arthropod species decreased slightly during intermediate stages (stage 2: β = −0.08, p < 0.001; stage 3: β = −0.05, p = 0.08; Model 7 - Table S1). In contrast, it increased markedly in the latest stage (stage 4: β = 0.43, p < 0.001, Model 7 - Table S1). Variance in trophic niche overlap also declined significantly with deglaciation stage, as indicated by the stage-dependent dispersion model, pointing to increasingly homogeneous dietary overlap patterns in later stages (stage 2: β = −0.13, p < 0.001; stage 3: β = −0.27, p = < 0.001; stage 4: β = −0.53, p = < 0.001; Model 7 – Table S1).

## DISCUSSION

Glacier forelands formed after glacier retreat represent real-world laboratories for studying the assembly and dynamics of species interaction networks over space and time. Although the impact of glacier retreat on biological communities including plants and insects has been well documented (Ficetola et al., 2021; Losapio et al., 2025a), the mechanisms underlying the development of plant–arthropod trophic interactions remain poorly understood. By using gut-content DNA metabarcoding techniques we detected the impact of glacier retreat on trophic niche overlap and niche breadth across multiple, diverse arthropod trophic guilds; as well as revealed otherwise cryptic diets. Moreover, we report that arthropod dietary composition diverges faster than changes in either plant or arthropod community composition following glacier retreat. Such rapid interaction reshuffling showed directional trends as trophic niche breadth decreased while trophic niche overlap increased with increasing age of deglaciation stages. Our findings suggest that trophic interaction networks are particularly key sensitive indicators of ecosystem response to climate change.

### Diet species richness declines but local diet increases following glacier retreat, reflecting the shift from pioneer communities to woody vegetation

Diet species richness showed a short-term increase followed by significant long-term decline with glacier extinction, confirming our first hypothesis that DSR decreases as succession approaches its late stages. Although plant species richness alone did not exert a significant main effect on DSR, its effect on arthropod diet richness was contingent on successional stage after glacier retreat, indicating that the relationship between plant richness and diet richness is not fixed but shifts as succession proceeds. Following glacier retreat, shifts in arthropod diet reflect both the changes in plant community composition (Moretti et al., 2006; Erschbamer & Caccianiga, 2016; Tu et al., 2024), dominance structure, food quality, also selective feeding by arthropod consumers (Scherber et al., 2010; Pellissier et al., 2012; Forister et al., 2015). Pioneer plants such as nitrogen-fixing Fabaceae and wind-dispersed Asteraceae were mostly consumed at early stages, while in the late deglaciation stage, the increasing prevalence of Ericaceae and Pinaceae in arthropod diets likely reflects both the structural dominance of these taxa in mature alpine ecosystems and the narrowing of available food choices as overall plant diversity declines (Vázquez et al. 2007). Guild-specific responses further underscore this complexity: pollinators tracked floral species diversity, while herbivores were associated to woody plant richness. Our findings highlight the importance of accounting for trophic guild identity when predicting how trophic networks will reorganize under global change, especially in terms of conservation and management: favouring one guild or one species could lead to the disadvantages of the others (Mouillot et al., 2013).

In contrast to DSR, local diet intensity – measured as the probability that a plant species present in a plot is also detected in arthropod diets - increases consistently following glacier retreat. First, this pattern of “the higher the plant abundance, the higher the consumption” confirms the costs for plants of being dominant, consistent with a consumer-mediated form of negative density dependence, whereby arthropods feeding pressure on locally abundant plants may slow competitive exclusion and buffer plant diversity, a key mechanism supporting species coexistence and maintaining high diversity (Keddy, 2017; Hülsmann et al., 2021). Second, this evidence suggests that arthropods increasingly track spatially available food resources as succession progresses (Root 1973, Vázquez et al., 2007; Xi et al., 2020; Pitteloud et al., 2023), rather than foraging randomly across the landscape. Importantly, this local dietary tracking was consistent across all trophic guilds as the positive relationship between local plant abundance and dietary detection did not differ between flying and non-flying arthropods, indicating that this pattern reflects food resource availability rather than dispersal ability. However, local plant abundance only partially explained diet composition, as arthropod diets were never entirely accounted for by plants present within the local plot, suggesting that a proportion of individuals consistently foraged beyond their immediate deglaciation stage and exploit diverse plant resources across the broader landscape matrix (Wiens 1976; Dunning et al., 1992). Taken together with our concurrent findings from trophic niche breadth (see below), the increase in local diet intensity at later stages therefore likely reflects constrained foraging choices as plant diversity declines, with specialist consumers concentrating on few locally available food resources (Morris, 2003; Costa-Pereira et al., 2019).

### Trophic turnover changes faster than species turnover following glacier retreat

We hypothesised that arthropod diet similarity would shift along with plant and arthropod community succession, as diet could reflect the food resources (plants) and feeding habit of consumers (arthropods). Interestingly, within-stage trophic similarity declines significantly following glacier retreat at faster rates than plant community similarity and arthropod community similarity. This faster increase in dietary heterogeneity relative to plant and arthropod community heterogeneity indicates that arthropods are actively partitioning food resources through behavioural mechanisms, not just passively tracking plant availability or simply reflecting species turnover. These results extend previous studies from different ecosystems showing that interaction network change correlates with species turnover rate (Poisot et al., 2012; CaraDonna et al., 2017). It is possible that stronger effects of steep environmental gradients and strong resource changes in glacial ecosystems force arthropods to rapidly shift and adapt their diet (Tylianakis et al., 2007; CaraDonna et al., 2017). Our study suggests that trophic interactions are not the simple reflection of community composition but rather driven by the complex interplay between food resource availability, consumer identity, and feeding selectivity (Poisot et al., 2015).

There are several mechanisms that contribute to explaining this phenomenon. First, even common arthropods distributed across all deglaciation stages can change their dietary preferences in response to changing food resources, leading to dietary differentiation despite taxonomic continuity (Bolnick et al., 2003; Des Roches et al., 2018) and consistent with the flexible feeding behaviour observed in glacier foreland arthropods (Hågvar et al., 2024). Second, the replacement of generalist arthropods at the early stages with more specialized species at the later stages can amplify dietary differences (Ficetola et al., 2021). Third, although we did not directly measure microhabitat heterogeneity along our study area, the shift of vegetation from pioneer herbs to woody species can generates increasingly differentiated along the chronosequence, allowing for greater dietary partitioning among coexisting consumer species even as plant diversity decreases (Tews et al., 2004).

This pattern has important implications for predicting trophic interaction networks’ response to environmental change: using only taxonomic, alpha or beta, diversity can underestimate the response of ecological systems. In fact, we found that arthropod composition similarity remained higher than trophic similarity, reflecting the shift from simple to complex trophic interactions in the later deglaciation stages, associated to potential higher ecosystem stability (Dunne et al., 2002; Gellner & McCann 2016) despite lower community heterogeneity.

### Low food resource diversity reduces trophic niche breadth and increases trophic niche overlap

In accordance with our third hypothesis, trophic niche breadth declined significantly at the latest deglaciation stage, indicating increased trophic specialization. Notably, this pattern varied among guilds: pollinators maintained broader diets compared to herbivores and omnivores, while other trophic groups showed no detectable differences. Similar patterns of decline trophic specialization under resource limitation have been documented globally as the resource diversity hypothesis, whereby consumers contract their diet range as the variety of available food decreases (Schoener 1974; Costa-Pereira et al., 2019). Similar patterns of trophic specialization under declining resource diversity have been documented globally in insect herbivores (Forister et al. 2015) and are consistent with succession-driven shifts from generalist to specialist species with post-glacial succession (Hanusch et al. 2022; Losapio et al. 2025a). Moreover, in the late deglaciation stage, the dominance of woody taxa from Ericaceae and Pinaceae which typically exhibit stronger and more complex phytochemical defences than pioneer species, may further shape dietary specialization by favouring arthropods with co-evolved counter-adaptations to specific plant defences over generalists (Coley et al. 1985; Futuyma and Agrawal 2009; Ali and Agrawal 2012).

Overall, in the whole study area, we observed exceptionally low trophic niche overlap, as only 17.61 % of the calculated niche overall values were higher than 0.30 (Grossman, 1986; Wei et al., 2025). This result suggests dietary specialization, likely to minimize competition between consumers in resource-poor area such as glacial ecosystems (Schoener 1974). Following glacier retreat, we observed a hump-shaped pattern on trophic niche overlap among arthropods: it decreased from early to intermediate successional stages but increased at the end of the succession. The initial decline is consistent with the classical competition theory of succession, which predicts that increasing species richness drives niche differentiation and reduces overlap (MacArthur & Levins, 1967; Schoener, 1974). However, the highest niche overlap at the latest deglaciation stage partially contrasts with classical predictions that stable communities should maintain low overlap through niche partitioning. Instead, this increase in niche overlap reflects the “resource concentration” effect: as plant diversity declines, few species dominate, multiple arthropods are forced to feed on the same food resources, increasing overlap (Root 1973; Schoener 1974; Costa-Pereira et al., 2019). This pattern has been documented in other stressed, resource-poor or disturbed environments such as high mountain lake following glacier retreat (Tiberti et al. 2020) and variable marine environments (Dehnhard et al. 2020). Our findings are coherent with a competition hump-shaped pattern along succession: few species, weak competition and high overlap early, peak diversity and competition, low overlap at intermediate stages, and resource-driven convergence late (Huston 1979; Schumann et al., 2016; Ficetola et al., 2021).

Taken together, these patterns indicate that late-successional, post-glacial communities may therefore be ecological sensitive due to the increasing dietary homogenization as multiple arthropod species converge on few dominant plant taxa, combined with declining diet species richness, narrows the functional diversity of trophic interactions and potentially reduces food web resilience in case of disturbance (Dunne et al. 2002; Gellner and McCann 2016)..

## Future directions

While our study provides comprehensive results on and novel insights into the assembly and development of plant–arthropod trophic interactions following glacier retreat, we also open several approaches for future research. First, future studies using quantitative diet data, rather than presence–absence of DNA markers would provide deeper insights into feeding intensity, resource preferences, and interaction strength, allowing a more precise assessment to study the important of rare food resources in shaping trophic networks. Second, integrating phytochemical data would allow testing the impact of biotic factors including plant chemical defences and nutritional quality on food web structure. Third, comparative studies across multiple biogeographic systems with different geohydrology, climate, and regional species pool would provide generality and context to our findings and reveal the extent to which the trophic patterns are universal features or be contingent on local conditions. Together, these directions would build on the value of moving beyond species lists by addressing species interactions and their networks for understanding biodiversity change and trophic responses to glacier retreat.

## Conclusions

Our findings support three key hypotheses about the ecological niche processes underlying trophic interactions assembly and food web dynamics following glacier retreat. First, diet species richness increases in the short term after glacier retreat but then decreases in the long term in mature forests, while local diet increases in the later deglaciation stage. Second, arthropod diet composition changes more rapidly than species composition. Third, we provide evidence for a “double squeeze”: trophic niche breadth declines while trophic niche overlap increases significantly with glacial extinction. This pattern indicates a phase transition from resource partitioning and potential for niche differentiation to homogeneous but specialized diet on few, abundant resources. Reconstructing trophic interaction networks will help not only to detect signals of biodiversity change but also to anticipate the responses of communities to climate change and glacier extinction.

## Acknowledgments*

This work was financially supported by the Swiss National Science Foundation (PZ00P3 202127), the Next Generation EU, Italian Ministry of Research and Education (PRIN 2022 PNRR P2022N5KYJ), and Biodiversa+ PrioritIce project (Project number: 31BD30_209629; grant agreement no. 101052342). We express our gratitude to the personnel at the Service des forêts, de la nature et du paysage of Canton Valais, Switzerland for granting sampling permit and for their support.

## Author Contributions

The author used Claude, version Sonnet 4.5, an AI-assisted tool, to improve spelling, grammar, clarity, and readability. After using this tool, the author carefully reviewed and edited the text and takes full responsibility for the final content.

BNT: Data curation, Formal analysis, Investigation, Methodology, Resources, Visualization, Writing – original draft, Writing – review & editing. EMC: Conceptualization, Investigation, Methodology, Writing – review & editing. JH: Methodology, Writing – review & editing. NK: Methodology, Writing – review & editing. MC: Conceptualization, Resources, Writing – review & editing. SR: Conceptualization, Resources, Writing – review & editing. LF: Conceptualization, Resources, Writing – review & editing. GL: Conceptualization, Funding acquisition, Project administration, Resources, Investigation, Supervision, Writing – original draft, Writing – review & editing.

## Data availability

Data and R scripts (no novel code) will be made publicly available on Zenodo upon manuscript acceptance.

